# miniBUSCO: a faster and more accurate reimplementation of BUSCO

**DOI:** 10.1101/2023.06.03.543588

**Authors:** Neng Huang, Heng Li

## Abstract

**Motivation:** Assembly completeness evaluation of genome assembly is a critical assessment of the accuracy and reliability of genomic data. An incomplete assembly can lead to errors in gene predictions, annotation, and other downstream analyses. BUSCO is one of the most widely used tools for assessing the completeness of genome assembly by comparing the presence of a set of single-copy orthologs conserved across a wide range of taxa. However, the runtime of BUSCO can be long, particularly for some large genome assemblies. It is a challenge for researchers to quickly iterate the genome assemblies or analyze a large number of assemblies.

**Results:** Here, we present miniBUSCO, an efficient tool for assessing the completeness of genome assemblies. miniBUSCO utilizes the protein-to-genome aligner miniprot and the datasets of conserved orthologous genes from BUSCO. Our evaluation of the real human assembly indicates that miniBUSCO achieves a 14-fold speedup over BUSCO. Furthermore, miniBUSCO reports a more accurate completeness of 99.6% than BUSCO’s completeness of 95.7%, which is in close agreement with the annotation completeness of 99.5% for T2T-CHM13.

**Availability:** https://github.com/huangnengCSU/minibusco.

**Supplementary information:** Supplementary data are available at *Bioinformatics* online.

## 1 Introduction

With the recent advances in sequencing technologies, especially long-read sequencing, genome assembly has undergone a revolution in terms of quality and completeness. Long-read sequencing technologies have improved genome assembly by enabling the generation of more complete and complex genome assemblies, including the telomere-to-telomere assemblies and haplotype-resolved assemblies, which were previously challenging to assemble using short-read sequencing (Wenger *et al*., 2019; Nurk *et al*., 2022). With the development of new assembly algorithms and improvements in computational resources, the time required to assemble a human genome has decreased significantly. It is now possible to assemble a human genome within 8 hours (Cheng *et al*., 2021, Cheng 2022).

Evaluating assembly completeness is a critical process that provides valuable information about the accuracy and reliability of the assembly and allows for finding potential errors such as missing or misassembled regions. Various tools have been developed for evaluating the completeness of genome assemblies. QUAST (Gurevich *et al*., 2013) is a widely used assembly quality evaluation tool by mapping the contigs/scaffolds of the assemblies to the reference genome and then reports the number of misassemblies, gaps, and various contiguity metrics including N50 and L50. For de novo assemblies, BUSCO (Benchmarking Universal Single-Copy Orthologs) is a popular tool for evaluating the completeness of assemblies (Simão *et al*., 2015; Manni *et al*., 2021). BUSCO utilizes a set of conserved single-copy orthologs that are expected to be present in the assemblies and employs gene predictors to confirm the presence of these genes in the assembly to assess the assembly completeness. However, BUSCO may underestimate completeness. For example, for the telomere-to-telomere (T2T) CHM13 assembly (Nurk *et al*., 2022), BUSCO reports a completeness of only 95.7%, but applying BUSCO to annotated protein-coding genes gives a completeness of 99.5%. In addition, BUSCO is inefficient. Assessing the completeness of a human genome assembly using BUSCO can take around 7 hours, which is close to the time required for assembling a human genome. With the increasing complexity and size of genome assemblies, there is an urgent need for more efficient and accurate completeness evaluation tools.

In this study, we will introduce miniBUSCO, a highly efficient tool for assessing assembly completeness based on the protein-to-genome aligner miniprot (Li, 2023) and BUSCO lineage datasets (Zdobnov *et al*., 2021). The evaluation of real datasets demonstrates that miniBUSCO has a significant speed-up than BUSCO while achieving higher accuracy.

## 2 Methods

First miniBUSCO downloads the corresponding BUSCO lineage dataset from https://busco-data.ezlab.org/v5/data/ to evaluate the input genome or assembly. Each lineage dataset contains several orthologous species with near-universally-distributed single-copy genes and each single-copy gene used to evaluate assembly completeness is present in at least 90% of the species. So, each single-copy gene usually has multiple protein sequences from different species (Simão *et al*., 2015). To evaluate the completeness of input assembly, we map the protein sequences of these single-copy genes to the assembly, followed by quantifying the occurrences of these single-copy genes in the assembly. In cases some segments are missing in the assembly, certain single-copy genes cannot be aligned. On the contrary, if the assembly contains falsely duplicated segments, some single-copy genes will be aligned to multiple positions in the assembly. Miniprot is a fast aligner for mapping protein sequences to the genome. Compared to similar tools, miniprot has the advantages of shorter runtime and accurate detection of splice junctions and frameshifts. MiniBUSCO performs one round of miniprot, while BUSCO performs two rounds of MetaEuk (Levy *et al*., 2020) with different parameters for high enough sensitivity.

Due to the similarity of sequences in the genome, even if some fragments are missing in the assembly, the protein sequences of the corresponding single-copy genes may be aligned to other paralogous positions in the assembly with lower identities. This can potentially lead to a decrease in the precision of assembly completeness assessment. Like BUSCO, we employ HMMER3 (Mistry *et al*., 2013) to confirm orthology. Only matches above the score cutoff defined in lineage files will be retained. Since each single-copy gene may have multiple protein sequences, we choose the protein sequence with the highest hmmersearch score to represent the single-copy gene.

After filtering paralogous gene matches, the remaining matches are categorized into one of four types: complete and single-copy, complete and duplicated, fragmented, or missing. A gene is considered missing if it has no alignment after HMMER filtration. It is fragmented if all its alignments are shorter than a length threshold defined by BUSCO. The rest of genes are considered complete. A complete gene is considered to have a single-copy in the assembly if it only has one alignment, or duplicated if it has multiple alignments. MiniBUSCO reports the proportion of genes falling into each of the four categories as the assessment of assembly completeness.

## 3 Results

To evaluate the performance of miniBUSCO, we compared it with BUSCO on various species, including seven reference genomes of model organisms: *Homo sapiens, Mus musculus, Arabidopsis thaliana, Zea mays, Drosophila melanogaster, Caenorhabditis elegans*, and *Saccharomyces cerevisiae*. Furthermore, we evaluated miniBUSCO and BUSCO on 81 PacBio HiFi assemblies of Metazoa and 22 PacBio HiFi assemblies of Viridiplantae obtained from the Darwin Tree of Life Project. To analyze the frameshifts reported by miniBUSCO, we ran miniBUSCO on several Nanopore assemblies and PacBio assemblies. The details of the datasets can be found in **Supplementary Table S1**. For BUSCO, we used the option ‘-m genome’ and the default gene predictor MetaEuk.

### 3.1 Evaluation of model organism reference genomes

Table 1 shows the comparison of miniBUSCO and BUSCO on the reference genomes of seven model organisms. For *A. thaliana* and *S. cerevisiae* reference, the results of miniBUSCO and BUSCO shows a difference of less than 1%. For *D. melanogaster, C. elegans* and *M. musculus*, the differences between the assessment results of miniBUSCO and BUSCO range from 1% to 3%. When evaluating the reference genomes of H. sapiens and Z. mays, miniBUSCO and BUSCO show a significant difference exceeding 3%. For most reference genomes, miniBUSCO reports higher completeness than BUSCO.

**Table 1.**
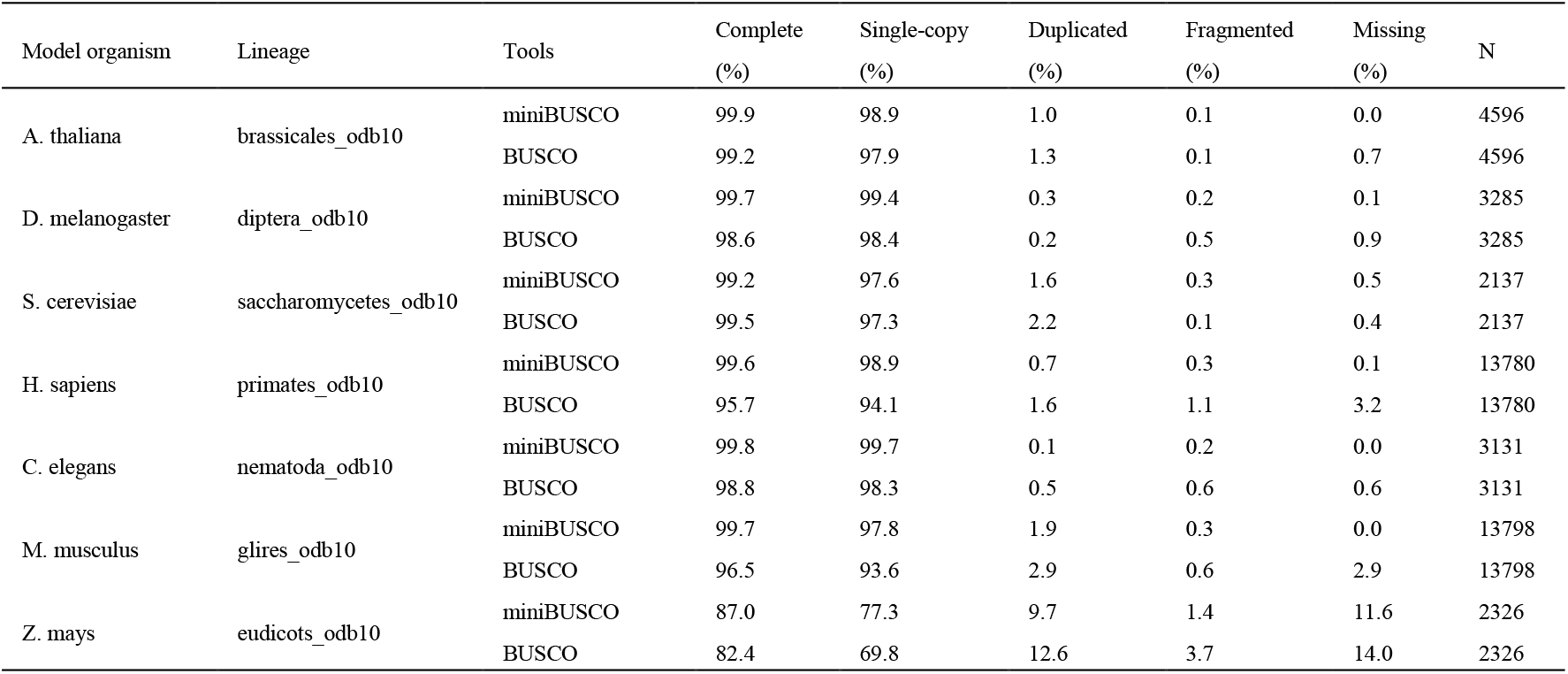
Comparison of miniBUSCO and BUSCO on the reference genomes of seven model organisms.

In the assessment of the *H. sapiens* reference genome (T2T-CHM13), miniBUSCO reported completeness of 99.6% compared to 95.7% reported by BUSCO. Among them, 562 complete genes were only reported by miniBUSCO. We subsequently evaluated these miniBUSCO-specific genes using the annotation of T2T-CHM13 from NCBI. By comparing the alignments of the complete genes with the annotation, we deemed a complete gene to be supported by annotation if the sum of overlap length in codons between the complete gene and annotation exceeds 50% of the total length of codons in annotation (see **Supplementary Figure S1**). We found that 557 out of 562 miniBUSCO-specific complete genes could be supported by the annotation.

Additionally, we used BUSCO protein mode to assess the completeness of T2T-CHM13 annotated gene set with the same lineage ‘primate_odb10’. We obtained the protein sequences of the T2T-CHM13 annotated gene set from NCBI and assessed the completeness with the command ‘*busco -m protein -i INPUT*.*amino_acids -o OUTPUT -l LINEAGE*’. In the protein mode, BUSCO searches the proteins of annotated gene set in the sequence database of the lineage dataset using HMMER3 (Mistry *et al*., 2013). **Supplementary Table S2** shows the assessment result of the annotated gene set, Complete: 99.5%, Fragmented: 0%, Missing: 0.5%. The completeness of annotated gene set is similar to the completeness of 99.6% obtained by miniBUSCO but differs from the completeness of 95.7% from BUSCO. By tracking the procedure of hmmsearch of these false negative complete genes in BUSCO, we observed that the protein sequences derived from the predicted genes by MetaEuk failed to meet the score thresholds. This indicates that these translated protein sequences have low quality, likely due to the errors in the protein-to-genome alignment. **Supplementary Figure S2** shows the protein-to-genome alignment of one of the miniBUSCO-specific complete genes. From that figure, we can see that some protein sequence fragments are missing in the MetaEuk alignment. While miniprot aligns the protein sequence almost completely to the CHM13 genome and the alignment is supported by the CHM13 annotation. The missing fragments by MetaEuk may be because MetaEuk cannot find the exact splice junction. For the *Z. mays* reference genome, miniBUSCO and BUSCO reported proportions of complete genes at 87.0% and 82.4%, respectively. We also evaluated the completeness of the annotation gene set of Z. mays (**Supplementary Table S3**) and the completeness of the annotation gene set is 87.2%, which is similar to the value obtained by miniBUSCO.

### 3.2 Evaluating the specificity of miniBUSCO

To check if miniBUSCO is overcalling complete genes, we removed chromosome 1 from T2T-CHM13 reference and compared the completeness of the modified CHM13 genome without chromosome 1 to the completeness of the entire CHM13 genome. **Supplementary Table S4** presents the assessment results by miniBUSCO. On the entire CHM13, miniBUSCO reported 13,622 complete single-copy genes, 1,435 of them were mapped to chromosome 1. After we removed chromosome 1 and ran miniBUSCO on the remaining chromosomes, miniBUSCO still reported 93 of them as complete single-copy genes. Checking the alignment of these 93 genes, we found that 60 of them were processed pseudogenes. As for the remaining 33 genes, their hits on other chromosomes were not as good as hits to chromosome 1 and suppressed by miniprot as miniprot was tuned to ignore alignments whose scores were below 97% of the best alignment scores. With chromosome 1 removed, the hits of genes to other chromosomes surfaced, passed the HMMER thresholds and became single-copy genes. Overall, the great majority of complete single-copy genes reported by miniBUSCO were real.

### 3.3 Evaluation of runtime

**Figure 1a** shows the runtimes of miniBUSCO and BUSCO on the seven reference genomes. All experiments were conducted on a single server with 40 threads. Notably, miniBUSCO achieves significant improvement over BUSCO by 3.4 to 14.5 times. Moreover, the speedup tends to increase with larger genome sizes, highlighting the efficiency of miniBUSCO in handling larger genomes. The evaluation of *H. sapiens* genome reduces from 6.7 hours to 0.4 hours. During the evaluation process of both miniBUSCO and BUSCO, the main time consumption is protein-to-genome alignment (MetaEuk/miniprot) and hmmsearch. There are two rounds of MetaEuk+hmmsearch in BUSCO while miniBUSCO only has one round of miniprot+hmmsearch. Meanwhile, miniprot has a 10-fold speedup over MetaEuk (Li, 2023). In summary, miniBUSCO can greatly improve the speed of evaluation. Furthermore, when dealing with small genomes, the evaluation process’s runtime is primarily influenced by hmmsearch, which relies on the size of the lineage dataset. However, for larger genomes, the running time of the evaluation process is mainly determined by protein-to-genome alignment. As the genome size increases, miniBUSCO demonstrates superior acceleration compared to BUSCO.

**Fig. 1.**
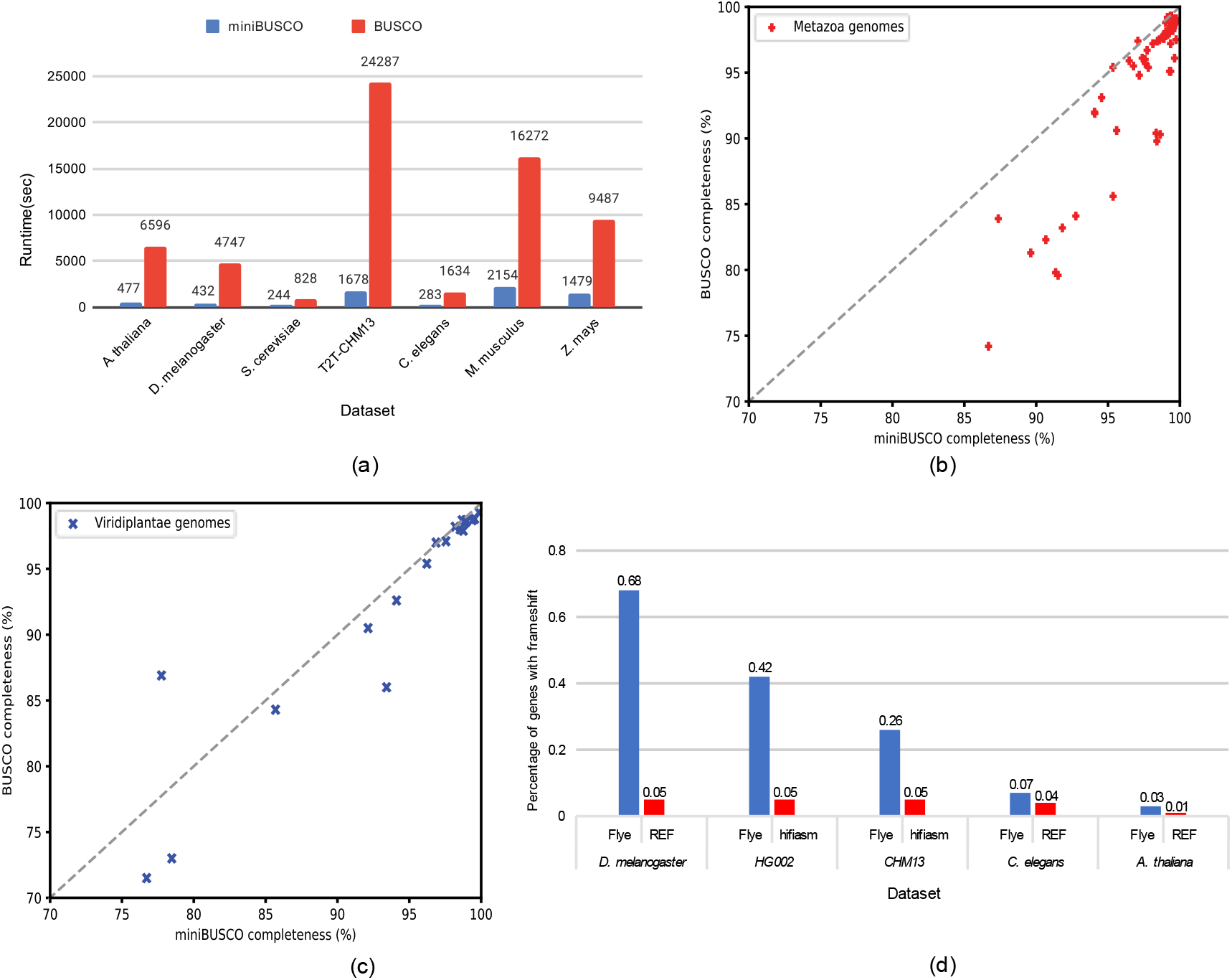
(a) Comparison of the runtime of miniBUSCO and BUSCO on real datasets. (b) The comparison of completeness reported by miniBUSCO and BUSCO on HiFi assemblies of Metazoa genomes. (c) The comparison of completeness reported by miniBUSCO and BUSCO on HiFi assemblies of Viridiplantae genomes. (d) The percentage of BUSCO genes with frameshifts reported by miniBUSCO on datasets of different species.

### 3.4 Evaluation of 103 HiFi assemblies of Metazoa and Viridiplantae

In addition to the reference genomes, we evaluated the completeness of 103 PacBio HiFi assemblies using miniBUSCO. The BUSCO assessments of these assemblies were downloaded from the website of Blobtoolkit2 (https://blobtoolkit.genomehubs.org) (Challis *et al*., 2020). **Supplementary Table S5** and **Supplementary Table S6** present the assessment results of 81 Metazoa assemblies and 22 Viridiplantae assemblies, respectively. **Figure 1b** shows the numbers of completed genes reported by miniBUSCO and BUSCO on HiFi assemblies of Metazoa genomes. On 78 out of 81 Metazoa assemblies, miniBUSCO reports a higher completeness range of 0-12% than BUSCO. For the remaining three Metazoa assemblies, there is a slight difference of less than 0.3% in completeness between miniBUSCO and BUSCO. **Figure 1c** presents the comparison of the completeness reported by miniBUSCO and BUSCO on HiFi assemblies of Viridiplantae genomes. On 20 out of 22 Viridiplantae assemblies, miniBUSCO obtained a higher completeness range of 0-7.5% than BUSCO. However, on the genome of *Pycnococcus provasolii*, miniBUSCO reported completeness that was 7.5% lower than that reported by BUSCO. By tracking the protein alignment of this genome, we found that most of the alignment records have very low identities due to the high divergence. Therefore, we add a feature to miniBUSCO that suggests running BUSCO additionally to ensure a more reliable assessment of assembly completeness for input assemblies with high divergence.

### 3.5 Frameshift analysis

Frameshift refers to the insertion or deletion of several base pairs that are not a multiple of three, disrupting the triplet reading frame of a DNA sequence. Miniprot can align through frameshifts in the genome sequence and identify them. MiniBUSCO outputs the number of frameshifts in each mapped gene by parsing the cigar sequences in the alignment output file. The number of frameshifts can be used as a measure of the quality of an assembly. We ran miniBUSCO and calculated the percentage of BUSCO genes with frameshifts on the assemblies or reference genomes of different species. We conducted a comparison of frameshifts in Flye assemblies and hifiasm assemblies for the genomes of HG002 and CHM13. The Flye assemblies of HG002 and CHM13 were assembled from 110×∼120× Nanopore reads by Flye (Kolmogorov *et al*., 2019) and the hifiasm assemblies of HG002 and CHM13 were generated from 32×∼36× PacBio HiFi reads by hifiasm (Cheng *et al*., 2021, 2022). **Figure 1d** shows the percentages of BUSCO genes with frameshifts in Flye assemblies of HG002 and CHM13 are 42% and 26%, respectively, whereas both hifiasm assemblies demonstrate a lower value of 5%. The Nanopore assemblies had a much higher frameshift error rate. We had a similar observation for *D. melanogaster, C. elegans*, and *A. thaliana*, assembled from 30× Nanopore reads, 40× PacBio CLR reads and 75× PacBio CLR reads, respectively. Nonetheless, the completeness assessments reported by miniBUSCO were not affected by the high error rate much (**Supplementary Table S7**). Separating frameshift errors from assembly incompleteness is a unique advantage of miniBUSCO.

## Discussion

This study shows that miniBUSCO is a highly efficient tool for evaluating the completeness of genome assemblies, offering improved accuracy and reduced evaluation time. However, it should be noted that miniBUSCO has limited sensitivity to distant homologs. Therefore, for assemblies with high divergence, combining the results from miniBUSCO and BUSCO is recommended to ensure higher reliability.

## Acknowledgements

We thank the Darwin Tree of Life project for releasing the data of Metazoa and Viridiplantae genomes.

## References

Wenger, A.M. et al. (2019) Accurate circular consensus long-read sequencing improves variant detection and assembly of a human genome. Nat. Biotechnol., 37, 1155–1162.

Nurk, S. et al. (2022) The complete sequence of a human genome. Science, 376, 44–53.

Cheng, H. et al. (2021) Haplotype-resolved de novo assembly using phased assembly graphs with hifiasm. Nat. methods, 18, 170–175.

Cheng, H. et al. (2022) Haplotype-resolved assembly of diploid genomes without parental data. Nat. Biotechnol., 40, 1332–1335.

Gurevich, A. et al. (2013) QUAST: quality assessment tool for genome assemblies. Bioinformatics, 29, 1072–1075.

Simão, F .A. et al. (2015) BUSCO: assessing genome assembly and annotation completeness with single-copy orthologs. Bioinformatics, 31, 3210–3212.

Manni, M. et al. (2021) BUSCO update: novel and streamlined workflows along with broader and deeper phylogenetic coverage for scoring of eukaryotic, prokaryotic, and viral genomes. Molecular biology and evolution, 38, 4647–4654.

Li, H. (2023) Protein-to-genome alignment with miniprot. Bioinformatics, Vol. 39, btad014.

Zdobnov, E. et al. (2021) OrthoDB in 2020: evolutionary and functional annotations of orthologs. Nucleic Acids Res., 49, D389–D393.

Levy, K. et al. (2020) MetaEuk-sensitive, high-throughput gene discovery, and annotation for large-scale eukaryotic metagenomics. Microbiome, 8, 1–15.

Mistry, J. et al. (2013) Challenges in homology search: HMMER3 and convergent evolution of coiled-coil regions. Nucleic Acids Res., 41, e121–e121.

Challis, R. et al. (2020) BlobToolKit–interactive quality assessment of genome assemblies. G3: Genes, Genomes, Genetics, 10, 1361–1374.

Kolmogorov, M. et al. (2019) Assembly of long, error-prone reads using repeat graphs. Nat. biotechnol., 37, 540–546.

