## Supplementary_material for "miniBUSCO: a faster and more accurate reimplementation of BUSCO"

### Supplementary Materials

#### Supplementary Tables

**Table S1.** The details of datasets.

| Dataset | Type | Accession |
|---|---|---|
| Arabidopsis thaliana | Reference | GCF_000001735.4_TAIR10.1_genomic.fna.gz |
| Drosophila melanogaster | Reference | GCF_000001215.4_Release_6_plus_ISO1_MT_genomic.fna.gz |
| Saccharomyces cerevisiae | Reference | GCF_000146045.2_R64_genomic.fna.gz |
| Homo sapiens | Reference | GCF_009914755.1_T2T-CHM13v2.0_genomic.fna.gz |
| Caenorhabditis elegans | Reference | GCF_000002985.6_WBcel235_genomic.fna.gz |
| Mus musculus | Reference | GCF_000001635.27_GRCm39/GCF_000001635.27_GRCm39_genomic.fna.gz |
| Zea mays | Reference | GCF_902167145.1_Zm-B73-REFERENCE-NAM-5.0_genomic.fna.gz |
| Metazoa | HiFi assemblies | https://portal.darwintreeoflife.org/data |
| Viridiplantae | HiFi assemblies | https://portal.darwintreeoflife.org/data |
| CHM13 | HiFi assembly | CHM13.HiFi.hifiasm-0.12.fa.gz |
| CHM13 | ONT assembly | flye.v29.chm13.ont.120x.fasta.gz |
| HG002 | HiFi assembly | NA24385.HiFi.hifiasm-0.12.pri.fa.gz |
| HG002 | ONT assembly | flye.v29.hg002.ont.110x.fasta.gz |
| Drosophila melanogaster | ONT assembly | flye.v29.dmelanogaster.ont.30x.fasta.gz |
| Caenorhabditis elegans | PacBio CLR assembly | flye.v29.celegans.pb.40x.fasta.gz |
| Arabidopsis thaliana | PacBio CLR assembly | flye.v29.athaliana.pb.75x.fasta.gz |

**Table S2.** The completeness of annotated gene set of T2T-CHM13 evaluated by BUSCO.

| C:99.5% [S:28.9%, D:70.6%], F:0.0%, M:0.5%, n:13780 | |
|---|---|
| Complete (C) | 13708 |
| Complete and single-copy (S) | 3976 |
| Complete and duplicated (D) | 9732 |
| Fragmented (F) | 4 |
| Missing (M) | 68 |
| Total (n) | 13780 |

**Table S3.** The completeness of annotated gene set of reference genome *Z. mays* evaluated by BUSCO.

| C:87.2% [S:46.7%, D:40.5%], F:0.8%, M:12.0%, n:2326 | |
|---|---|
| Complete (C) | 2028 |
| Complete and single-copy (S) | 1086 |
| Complete and duplicated (D) | 942 |
| Fragmented (F) | 18 |
| Missing (M) | 280 |
| Total (n) | 2326 |

**Table S4.** The assessment of completeness of the entire T2T-CHM13 genome and the modified T2T-CHM13 without chromosome 1.

| Dataset | Completed (%) | Single-copy (%) | Duplicated (%) | Fragmented (%) | Missing (%) | N |
|---|---|---|---|---|---|---|
| T2T-CHM13 | 99.6 | 98.9 | 0.7 | 0.3 | 0.1 | 13780 |
| T2T-CHM13-without-chr1 | 89.9 | 89.2 | 0.7 | 0.4 | 9.7 | 13780 |

**Table S5.** The assessment of completeness of 81 HiFi assemblies of Metazoa by miniBUSCO and BUSCO

| Accession | Taxon | Lineage | miniBUSCO | | | | | BUSCO | | | | |
|---|---|---|---|---|---|---|---|---|---|---|---|---|
| | | | complete | single | duplicated | fragmented | missing | complete | single | duplicated | fragmented | missing |
| GCA_943735955.1 | Piscicola geometra | metazoa | 0.867 | 0.862 | 0.005 | 0.018 | 0.115 | 0.742 | 0.735 | 0.007 | 0.090 | 0.168 |
| GCA_922989275.1 | Gari tellinella | mollusca | 0.915 | 0.906 | 0.009 | 0.010 | 0.074 | 0.796 | 0.784 | 0.012 | 0.048 | 0.157 |
| GCA_947247005.1 | Spisula solida | mollusca | 0.913 | 0.905 | 0.008 | 0.013 | 0.075 | 0.798 | 0.787 | 0.011 | 0.046 | 0.156 |
| GCA_921293015.1 | Phorcus lineatus | mollusca | 0.954 | 0.949 | 0.005 | 0.012 | 0.035 | 0.856 | 0.850 | 0.006 | 0.044 | 0.100 |
| GCA_935421135.1 | Bugulina stolonifera | metazoa | 0.928 | 0.925 | 0.003 | 0.013 | 0.060 | 0.841 | 0.831 | 0.009 | 0.070 | 0.089 |
| GCA_945261195.1 | Cryptosula pallasiana | metazoa | 0.918 | 0.915 | 0.003 | 0.018 | 0.064 | 0.832 | 0.821 | 0.012 | 0.080 | 0.088 |
| GCA_929108145.1 | Limnephilus rhombicus | endopterygota | 0.984 | 0.976 | 0.008 | 0.008 | 0.009 | 0.898 | 0.886 | 0.012 | 0.071 | 0.032 |
| GCA_936435175.1 | Glyphotaelius pellucidus | endopterygota | 0.986 | 0.981 | 0.006 | 0.007 | 0.007 | 0.903 | 0.895 | 0.008 | 0.068 | 0.029 |
| GCA_916610825.1 | Haliclystus octoradiatus | metazoa | 0.896 | 0.894 | 0.002 | 0.017 | 0.087 | 0.813 | 0.811 | 0.002 | 0.094 | 0.092 |
| GCA_914767715.1 | Membranipora membranacea | metazoa | 0.906 | 0.893 | 0.013 | 0.019 | 0.076 | 0.823 | 0.799 | 0.024 | 0.083 | 0.094 |
| GCA_917880885.1 | Limnephilus marmoratus | endopterygota | 0.984 | 0.977 | 0.007 | 0.008 | 0.009 | 0.904 | 0.893 | 0.011 | 0.067 | 0.029 |
| GCA_945859605.1 | Lumbricus rubellus | metazoa | 0.956 | 0.937 | 0.019 | 0.009 | 0.035 | 0.906 | 0.869 | 0.037 | 0.046 | 0.048 |
| GCA_922984935.1 | Meles meles | carnivora | 0.994 | 0.976 | 0.018 | 0.004 | 0.003 | 0.951 | 0.932 | 0.020 | 0.008 | 0.040 |
| GCA_922990625.1 | Meles meles | carnivora | 0.993 | 0.977 | 0.016 | 0.004 | 0.003 | 0.951 | 0.930 | 0.021 | 0.008 | 0.041 |
| GCA_910594005.1 | Cervus elaphus | cetartiodactyla | 0.996 | 0.971 | 0.025 | 0.003 | 0.001 | 0.961 | 0.928 | 0.033 | 0.011 | 0.028 |
| GCA_916048095.1 | Sacculina carcini | arthropoda | 0.874 | 0.870 | 0.004 | 0.014 | 0.113 | 0.839 | 0.832 | 0.007 | 0.061 | 0.100 |
| GCA_942159475.1 | Sthenelais limicola | metazoa | 0.978 | 0.970 | 0.008 | 0.006 | 0.016 | 0.954 | 0.943 | 0.010 | 0.028 | 0.018 |
| GCA_932526225.1 | Nematostella vectensis | metazoa | 0.972 | 0.969 | 0.003 | 0.004 | 0.024 | 0.948 | 0.942 | 0.005 | 0.025 | 0.027 |
| GCA_929443795.1 | Accipiter gentilis | aves | 0.997 | 0.994 | 0.003 | 0.002 | 0.001 | 0.975 | 0.967 | 0.008 | 0.006 | 0.019 |
| GCA_946894095.1 | Incurvaria masculella | lepidoptera | 0.941 | 0.937 | 0.004 | 0.007 | 0.053 | 0.919 | 0.911 | 0.008 | 0.014 | 0.067 |
| GCA_921293095.1 | Ischnura elegans | insecta | 0.993 | 0.987 | 0.006 | 0.003 | 0.004 | 0.972 | 0.964 | 0.008 | 0.013 | 0.015 |
| GCA_946902875.1 | Nematopogon swammerdamellus | lepidoptera | 0.941 | 0.935 | 0.005 | 0.007 | 0.053 | 0.920 | 0.909 | 0.011 | 0.013 | 0.067 |
| GCA_936440205.1 | Lepidonotus clava | metazoa | 0.976 | 0.974 | 0.002 | 0.006 | 0.018 | 0.957 | 0.953 | 0.004 | 0.023 | 0.020 |
| GCA_918808275.1 | Dolichovespula sylvestris | hymenoptera | 0.976 | 0.959 | 0.017 | 0.009 | 0.016 | 0.960 | 0.957 | 0.003 | 0.010 | 0.030 |
| GCA_918807975.1 | Aplidium turbinatum | metazoa | 0.946 | 0.921 | 0.024 | 0.013 | 0.042 | 0.931 | 0.888 | 0.043 | 0.034 | 0.036 |
| GCA_947063395.1 | Hemistola chrysoprasaria | lepidoptera | 0.995 | 0.992 | 0.003 | 0.003 | 0.002 | 0.982 | 0.978 | 0.004 | 0.004 | 0.013 |
| GCA_947310845.1 | Yponomeuta plumbellus | lepidoptera | 0.989 | 0.987 | 0.002 | 0.003 | 0.008 | 0.976 | 0.972 | 0.004 | 0.007 | 0.017 |
| GCA_918843875.1 | Diadumene lineata | metazoa | 0.974 | 0.972 | 0.002 | 0.007 | 0.019 | 0.961 | 0.956 | 0.005 | 0.020 | 0.019 |
| GCA_947458855.1 | Monopis laevigella | lepidoptera | 0.968 | 0.962 | 0.006 | 0.006 | 0.027 | 0.955 | 0.946 | 0.009 | 0.008 | 0.037 |
| GCA_923062675.1 | Marasmarcha lunaedactyla | lepidoptera | 0.988 | 0.982 | 0.006 | 0.003 | 0.008 | 0.976 | 0.967 | 0.009 | 0.007 | 0.017 |
| GCA_934047225.1 | Ypsolopha sequella | lepidoptera | 0.992 | 0.985 | 0.007 | 0.003 | 0.005 | 0.980 | 0.969 | 0.012 | 0.005 | 0.015 |
| GCA_947086465.1 | Eupithecia exiguata | lepidoptera | 0.991 | 0.988 | 0.003 | 0.003 | 0.006 | 0.979 | 0.974 | 0.005 | 0.006 | 0.015 |
| GCA_947172395.1 | Bicyclus anynana | lepidoptera | 0.994 | 0.989 | 0.005 | 0.003 | 0.003 | 0.982 | 0.976 | 0.006 | 0.004 | 0.014 |
| GCA_947049275.1 | Epinotia bilunana | lepidoptera | 0.991 | 0.988 | 0.003 | 0.003 | 0.006 | 0.979 | 0.974 | 0.005 | 0.006 | 0.016 |
| GCA_934044485.1 | Apeira syringaria | lepidoptera | 0.996 | 0.990 | 0.006 | 0.003 | 0.002 | 0.984 | 0.977 | 0.007 | 0.005 | 0.010 |
| GCA_947507515.1 | Eulithis testata | lepidoptera | 0.993 | 0.992 | 0.001 | 0.003 | 0.004 | 0.981 | 0.978 | 0.004 | 0.005 | 0.014 |
| GCA_947044415.1 | Eupithecia dodoneata | lepidoptera | 0.991 | 0.989 | 0.003 | 0.003 | 0.005 | 0.980 | 0.976 | 0.004 | 0.005 | 0.014 |
| GCA_932527185.1 | Ecliptopera silaceata | lepidoptera | 0.994 | 0.991 | 0.003 | 0.003 | 0.003 | 0.983 | 0.979 | 0.004 | 0.004 | 0.013 |
| GCA_910589615.1 | Taurulus bubalis | actinopterygii | 0.995 | 0.991 | 0.004 | 0.004 | 0.002 | 0.984 | 0.976 | 0.008 | 0.005 | 0.011 |
| GCA_934045075.1 | Yponomeuta sedellus | lepidoptera | 0.986 | 0.982 | 0.004 | 0.003 | 0.011 | 0.975 | 0.970 | 0.005 | 0.006 | 0.019 |
| GCA_933210815.1 | Meta bourneti | arachnida | 0.990 | 0.956 | 0.033 | 0.002 | 0.008 | 0.979 | 0.922 | 0.057 | 0.008 | 0.013 |
| GCA_947359385.1 | Chesias legatella | lepidoptera | 0.995 | 0.992 | 0.003 | 0.002 | 0.003 | 0.984 | 0.980 | 0.004 | 0.005 | 0.012 |
| GCA_921293045.1 | Aplocera efformata | lepidoptera | 0.995 | 0.993 | 0.002 | 0.003 | 0.003 | 0.984 | 0.981 | 0.003 | 0.004 | 0.012 |
| GCA_947310995.1 | Yponomeuta cagnagella | lepidoptera | 0.988 | 0.986 | 0.003 | 0.004 | 0.008 | 0.978 | 0.974 | 0.004 | 0.005 | 0.018 |
| GCA_932526625.1 | Rhingia campestris | diptera | 0.984 | 0.975 | 0.009 | 0.005 | 0.011 | 0.974 | 0.959 | 0.014 | 0.008 | 0.018 |
| GCA_946894065.1 | Acleris literana | lepidoptera | 0.991 | 0.988 | 0.004 | 0.003 | 0.006 | 0.981 | 0.974 | 0.007 | 0.004 | 0.015 |
| GCA_945859705.1 | Episyrphus balteatus | diptera | 0.977 | 0.973 | 0.005 | 0.007 | 0.016 | 0.967 | 0.960 | 0.007 | 0.007 | 0.026 |
| GCA_923062465.1 | Acleris sparsana | lepidoptera | 0.993 | 0.990 | 0.004 | 0.002 | 0.005 | 0.983 | 0.977 | 0.006 | 0.004 | 0.013 |
| GCA_947284805.1 | Coleophora flavipennella | lepidoptera | 0.989 | 0.982 | 0.008 | 0.003 | 0.008 | 0.979 | 0.968 | 0.011 | 0.005 | 0.017 |
| GCA_947369235.1 | Udea olivalis | lepidoptera | 0.996 | 0.993 | 0.004 | 0.003 | 0.001 | 0.987 | 0.984 | 0.004 | 0.003 | 0.010 |
| GCA_905147045.1 | Nymphalis io | lepidoptera | 0.997 | 0.993 | 0.004 | 0.003 | 0.000 | 0.988 | 0.985 | 0.003 | 0.004 | 0.009 |
| GCA_947086405.1 | Esperia sulphurella | lepidoptera | 0.990 | 0.985 | 0.005 | 0.003 | 0.007 | 0.981 | 0.975 | 0.007 | 0.003 | 0.016 |
| GCA_946965025.1 | Blera fallax | diptera | 0.981 | 0.978 | 0.004 | 0.006 | 0.013 | 0.972 | 0.968 | 0.004 | 0.008 | 0.020 |
| GCA_922987775.1 | Agonopterix subpropinquella | lepidoptera | 0.996 | 0.990 | 0.007 | 0.002 | 0.002 | 0.987 | 0.979 | 0.007 | 0.002 | 0.012 |
| GCA_947044815.1 | Apomyelois bistriatella | lepidoptera | 0.997 | 0.994 | 0.003 | 0.003 | 0.000 | 0.988 | 0.983 | 0.005 | 0.003 | 0.009 |
| GCA_947034925.1 | Eudemis profundana | lepidoptera | 0.994 | 0.987 | 0.007 | 0.003 | 0.004 | 0.985 | 0.974 | 0.011 | 0.004 | 0.011 |
| GCA_921293315.1 | Nemurella pictetii | insecta | 0.996 | 0.990 | 0.007 | 0.002 | 0.002 | 0.988 | 0.980 | 0.008 | 0.004 | 0.008 |
| GCA_947508005.1 | Amphipoea lucens | lepidoptera | 0.995 | 0.992 | 0.003 | 0.002 | 0.003 | 0.987 | 0.981 | 0.006 | 0.003 | 0.010 |
| GCA_945859575.1 | Herminia tarsipennalis | lepidoptera | 0.994 | 0.987 | 0.007 | 0.002 | 0.003 | 0.986 | 0.977 | 0.009 | 0.003 | 0.011 |
| GCA_905146925.1 | Autographa gamma | lepidoptera | 0.996 | 0.994 | 0.003 | 0.003 | 0.001 | 0.988 | 0.986 | 0.003 | 0.002 | 0.009 |
| GCA_932526445.1 | Phragmatobia fuliginosa | lepidoptera | 0.995 | 0.989 | 0.006 | 0.003 | 0.002 | 0.987 | 0.979 | 0.008 | 0.003 | 0.010 |
| GCA_936440315.1 | Barbus barbus | actinopterygii | 0.995 | 0.205 | 0.790 | 0.004 | 0.001 | 0.987 | 0.210 | 0.776 | 0.005 | 0.008 |
| GCA_927399405.1 | Agonopterix arenella | lepidoptera | 0.995 | 0.990 | 0.005 | 0.002 | 0.003 | 0.987 | 0.980 | 0.007 | 0.003 | 0.010 |
| GCA_947462355.1 | Caradrina kadenii | lepidoptera | 0.996 | 0.995 | 0.001 | 0.003 | 0.001 | 0.989 | 0.986 | 0.003 | 0.003 | 0.008 |
| GCA_921972225.1 | Euplexia lucipara | lepidoptera | 0.997 | 0.995 | 0.003 | 0.002 | 0.000 | 0.990 | 0.985 | 0.005 | 0.003 | 0.007 |
| GCA_918843915.1 | Notodonta ziczac | lepidoptera | 0.996 | 0.992 | 0.004 | 0.003 | 0.002 | 0.989 | 0.984 | 0.004 | 0.003 | 0.009 |
| GCA_947347705.1 | Nymphula nitidulata | lepidoptera | 0.996 | 0.994 | 0.002 | 0.003 | 0.002 | 0.989 | 0.986 | 0.003 | 0.002 | 0.009 |
| GCA_934045905.1 | Leuctra nigra | insecta | 0.996 | 0.980 | 0.016 | 0.002 | 0.003 | 0.989 | 0.970 | 0.019 | 0.005 | 0.006 |
| GCA_947256265.1 | Acronicta leporina | lepidoptera | 0.998 | 0.996 | 0.001 | 0.002 | 0.000 | 0.991 | 0.988 | 0.003 | 0.002 | 0.007 |
| GCA_947361185.1 | Charanyca ferruginea | lepidoptera | 0.995 | 0.967 | 0.028 | 0.002 | 0.002 | 0.989 | 0.959 | 0.030 | 0.002 | 0.009 |
| GCA_947363495.1 | Euzophera pinguis | lepidoptera | 0.995 | 0.992 | 0.003 | 0.003 | 0.002 | 0.989 | 0.985 | 0.003 | 0.003 | 0.008 |
| GCA_916618015.1 | Xestia c-nigrum | lepidoptera | 0.994 | 0.990 | 0.004 | 0.003 | 0.003 | 0.988 | 0.982 | 0.005 | 0.002 | 0.010 |
| GCA_921293005.1 | Nemoura dubitans | insecta | 0.994 | 0.985 | 0.009 | 0.002 | 0.004 | 0.988 | 0.977 | 0.012 | 0.004 | 0.007 |
| GCA_905475395.1 | Chrysoperla carnea | endopterygota | 0.964 | 0.960 | 0.005 | 0.008 | 0.028 | 0.959 | 0.950 | 0.009 | 0.010 | 0.031 |
| GCA_944039245.1 | Nebria salina | endopterygota | 0.990 | 0.989 | 0.001 | 0.007 | 0.003 | 0.988 | 0.984 | 0.003 | 0.007 | 0.005 |
| GCA_933228885.1 | Leistus spinibarbis | endopterygota | 0.993 | 0.992 | 0.001 | 0.005 | 0.001 | 0.992 | 0.987 | 0.004 | 0.005 | 0.003 |
| GCA_933228675.1 | Stomorhina lunata | diptera | 0.994 | 0.990 | 0.005 | 0.004 | 0.002 | 0.993 | 0.988 | 0.005 | 0.002 | 0.005 |
| GCA_911728435.1 | Apoderus coryli | endopterygota | 0.993 | 0.983 | 0.010 | 0.004 | 0.002 | 0.993 | 0.977 | 0.016 | 0.004 | 0.003 |
| GCA_922984085.1 | Sicus ferrugineus | diptera | 0.953 | 0.948 | 0.006 | 0.010 | 0.037 | 0.954 | 0.943 | 0.010 | 0.007 | 0.039 |
| GCA_930367205.1 | Acanthosoma haemorrhoidale | hemiptera | 0.991 | 0.981 | 0.010 | 0.001 | 0.008 | 0.992 | 0.974 | 0.018 | 0.001 | 0.007 |
| GCA_914767665.1 | Harmonia axyridis | endopterygota | 0.971 | 0.957 | 0.014 | 0.008 | 0.022 | 0.974 | 0.952 | 0.023 | 0.006 | 0.020 |

**Table S6.** The assessment of completeness of 22 HiFi assemblies of Viridiplantae by miniBUSCO and BUSCO

| Accession | Taxon | Lineage | miniBUSCO | | | | | BUSCO | | | | |
|---|---|---|---|---|---|---|---|---|---|---|---|---|
| | | | complete | single | duplicated | fragmented | missing | complete | single | duplicated | fragmented | missing |
| GCA_914767535.1 | Dunaliella primolecta | chlorophyta | 0.934 | 0.929 | 0.005 | 0.003 | 0.063 | 0.860 | 0.854 | 0.006 | 0.040 | 0.100 |
| GCA_946800655.1 | Juncus effusus | poales | 0.785 | 0.754 | 0.031 | 0.018 | 0.198 | 0.730 | 0.680 | 0.050 | 0.030 | 0.240 |
| GCA_946800325.1 | Luzula sylvatica | poales | 0.767 | 0.725 | 0.042 | 0.017 | 0.216 | 0.715 | 0.664 | 0.051 | 0.028 | 0.257 |
| GCA_933775445.1 | Potentilla anserina | eudicots | 0.921 | 0.911 | 0.011 | 0.015 | 0.064 | 0.905 | 0.888 | 0.017 | 0.017 | 0.079 |
| GCA_934048045.1 | Polygonum aviculare | eudicots | 0.941 | 0.902 | 0.040 | 0.008 | 0.051 | 0.926 | 0.869 | 0.057 | 0.018 | 0.056 |
| GCA_947034895.1 | Rhytidiadelphus loreus | embryophyta | 0.856 | 0.809 | 0.047 | 0.034 | 0.110 | 0.843 | 0.776 | 0.067 | 0.029 | 0.128 |
| GCA_947034885.1 | Thuidium tamariscinum | embryophyta | 0.618 | 0.587 | 0.031 | 0.034 | 0.348 | 0.605 | 0.563 | 0.042 | 0.027 | 0.367 |
| GCA_937616625.1 | Mercurialis annua | eudicots | 0.988 | 0.974 | 0.014 | 0.003 | 0.009 | 0.979 | 0.959 | 0.020 | 0.006 | 0.016 |
| GCA_946814005.1 | Chamaenerion angustifolium | eudicots | 0.962 | 0.828 | 0.135 | 0.008 | 0.030 | 0.954 | 0.796 | 0.158 | 0.011 | 0.035 |
| GCA_946800305.1 | Medicago arabica | fabales | 0.996 | 0.980 | 0.016 | 0.002 | 0.003 | 0.988 | 0.966 | 0.022 | 0.002 | 0.010 |
| GCA_946807835.1 | Ailanthus altissimus | eudicots | 0.994 | 0.613 | 0.381 | 0.003 | 0.003 | 0.987 | 0.547 | 0.440 | 0.003 | 0.009 |
| GCA_933208065.1 | Arabidopsis thaliana | brassicales | 0.998 | 0.988 | 0.010 | 0.001 | 0.000 | 0.993 | 0.979 | 0.014 | 0.001 | 0.007 |
| GCA_946800315.1 | Misopates orontium | eudicots | 0.985 | 0.961 | 0.025 | 0.004 | 0.010 | 0.980 | 0.948 | 0.032 | 0.003 | 0.016 |
| GCA_948329865.1 | Linaria vulgaris | eudicots | 0.976 | 0.944 | 0.032 | 0.007 | 0.017 | 0.971 | 0.926 | 0.045 | 0.006 | 0.023 |
| GCA_916050505.1 | Malus domestica | eudicots | 0.989 | 0.632 | 0.356 | 0.004 | 0.007 | 0.985 | 0.595 | 0.390 | 0.005 | 0.011 |
| GCA_916048215.1 | Malus sylvestris | eudicots | 0.989 | 0.628 | 0.361 | 0.004 | 0.007 | 0.985 | 0.594 | 0.391 | 0.006 | 0.009 |
| GCA_916615385.1 | Malus domestica | eudicots | 0.990 | 0.629 | 0.361 | 0.004 | 0.006 | 0.986 | 0.595 | 0.391 | 0.006 | 0.008 |
| GCA_916612005.1 | Malus domestica | eudicots | 0.989 | 0.625 | 0.364 | 0.004 | 0.007 | 0.987 | 0.599 | 0.388 | 0.005 | 0.008 |
| GCA_946896805.1 | Scutellaria galericulata | eudicots | 0.982 | 0.948 | 0.034 | 0.005 | 0.013 | 0.982 | 0.939 | 0.043 | 0.002 | 0.016 |
| GCA_946800695.1 | Geum urbanum | eudicots | 0.987 | 0.254 | 0.733 | 0.004 | 0.009 | 0.987 | 0.186 | 0.801 | 0.003 | 0.011 |
| GCA_947034835.1 | Ballota nigra | eudicots | 0.969 | 0.942 | 0.027 | 0.008 | 0.024 | 0.970 | 0.929 | 0.040 | 0.003 | 0.028 |
| GCA_938743325.1 | Pycnococcus provasolii | chlorophyta | 0.778 | 0.764 | 0.014 | 0.007 | 0.216 | 0.869 | 0.837 | 0.032 | 0.011 | 0.120 |

**Table S7.** The assessment of completeness of several datasets of Flye assemblies, hifiasm assemblies and reference genomes by miniBUSCO

| Dataset | Lineage | Completed (%) | Single-copy (%) | Duplicated (%) | Fragmented (%) | Missing (%) | N |
|---|---|---|---|---|---|---|---|
| HG002-Flye | primate_odb10 | 99.4 | 98.7 | 0.7 | 0.4 | 0.2 | 13780 |
| HG002-hifiasm | primate_odb10 | 99.5 | 98.7 | 0.8 | 0.4 | 0.2 | 13780 |
| CHM13-Flye | primate_odb10 | 99.5 | 98.8 | 0.7 | 0.4 | 0.2 | 13780 |
| CHM13-hifiasm | primate_odb10 | 99.5 | 98.8 | 0.7 | 0.4 | 0.1 | 13780 |
| A. thaliana-Flye | brassicales_odb10 | 99.8 | 98.8 | 1.1 | 0.1 | 0.0 | 4596 |
| A. thaliana-reference | brassicales_odb10 | 99.9 | 98.9 | 1.0 | 0.1 | 0.0 | 4596 |
| C. elegans-Flye | nematoda_odb10 | 99.8 | 99.6 | 0.2 | 0.2 | 0.0 | 3131 |
| C. elegans-reference | nematoda_odb10 | 99.8 | 99.7 | 0.1 | 0.2 | 0.0 | 3131 |
| D. melanogaster-Flye | diptera_odb10 | 99.6 | 99.5 | 0.1 | 0.3 | 0.1 | 3285 |
| D. melanogaster-reference | diptera_odb10 | 99.7 | 99.4 | 0.3 | 0.2 | 0.1 | 3285 |

**Supplementary Figures**

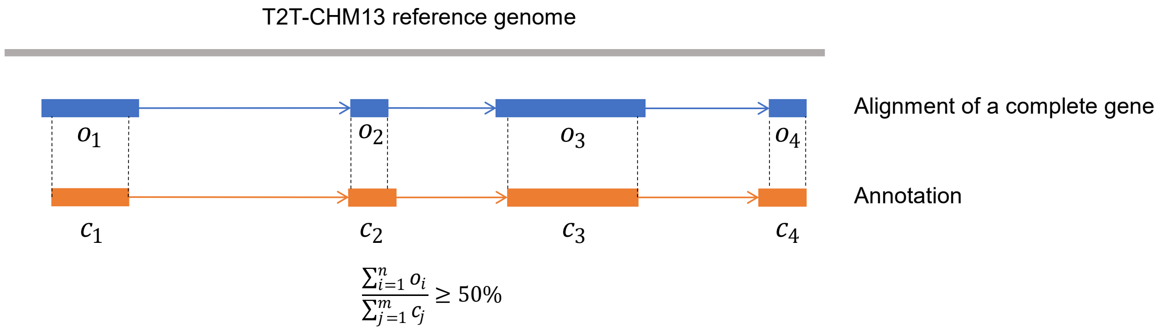


**Figure S1.** Example of the complete gene confirmed by the annotation. $o_i$ is the length of overlaps in codons between complete gene and annotation. $c_j$ is the length of a codon in the annotation.

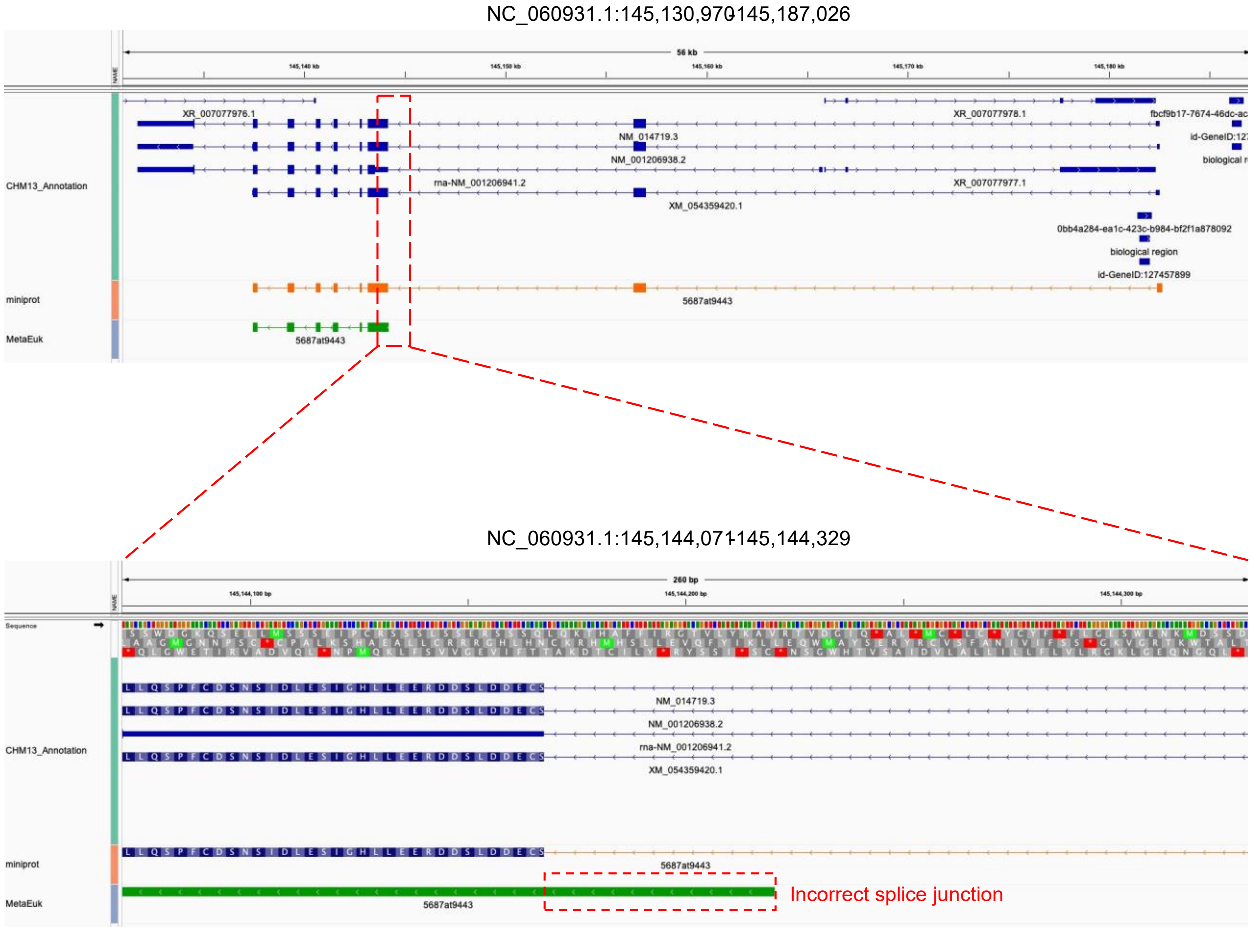


**Figure S2.** Example of the protein-to-genome alignment of a miniBUSCO-specific complete gene. The upper part of the figure shows that miniprot aligns the protein sequence almost completely to the CHM13 genome, and is supported by the CHM13 annotation, while MetaEuk misses some fragments when aligning the protein sequence. The lower part of the figure shows that the reason why some fragments are missing during the MetaEuk alignment may be that MetaEuk cannot find exact splice junctions.
